# Raltitrexed and Edatrexate drug molecules against human Proton Coupled Folate Transporter to cure cancer

**DOI:** 10.1101/2022.07.19.500571

**Authors:** Yashpal Singh Raja, Surya Mishra, Shikha Bhushan

## Abstract

Proton pump inhibitors (PPIs) (e.g.: - rabeprazole, pantoprazole, omeprazole, lansoprazole, and esomeprazole) are widely used to treat gastroesophageal reflux disease and other acid-related disorders. Folic acid transport is important for proper cell proliferation. In human, folic acid is not synthesized in the body, it is obtained externally. Therefore, specific transporters are involved in absorption of folic acid, which is concentrated in intestine. Malignant cancer cells require this folic acid very frequently in large quantities for the rapid reproduction of cancer cell. In the current study, we compared different types of proton pump inhibitor drug molecules that may be potential candidates for preventing the absorption of folate by (hPCFT), its responsible for unusually large and frequent folate production. Each of these anti-drug candidates has 27 finalists based on previous studies. Among these competitors, leukovorin and noletraxid molecules were found to be particularly associated with the hPCFT transporter’s key active site loop(G^155^XXG^158^).

## INTRODUCTION

In higher organisms, the absence of folate synthesizing protein indicates the presence of multiple folate transporters by protecting exogenous folate between cells and compartments such as mitochondria, cytoplasm, plastids, and vacuoles. Because folic acid is a hydrophilic molecule, it does not diffuse through biological membranes. Thus, in mammals, placental cells have adapted to a healthy transport system for cellular uptake of folate cofactors.

Genetic molecular and biological investigations reported two transporter families for classification of transporter i.e., ABC belong to general primary active transporters and are introduced in transportation of primary active macromolecules whereas MFS family functions in transportation of secondary active macromolecule in response to chemo-osmotic ion gradient.

MFS family of transporter are involved in transport of various collections of substrates like sugars, amino acids, nucleosides, sugar, organic phosphates and vitamins in antiport, symport and uniport manner. It is among the largest active secondary transporter family found in bacteria, archaebacteria and eukaryotes. The basic fold of MFS protein is 12 transmembrane helices with two 6 helixes bundles made up of N and C terminal homologous domain. To figure out the mechanism of membrane transport, it is important to analyse the protein structural information. Accordingly, structural predictions have relied on comprehensive array of biophysical and biochemical approaches along with structural modelling. (Hanson, A.D. and Roje, Folic acids and folates: the feasibility for nutritional enhancement in plant foods. Journal of Science of foods and Agriculture, 80(7), pp.795-824, 2000.)

There are basically three types of folate transporter: (RFC), (PCFT) and (FRs). It is located on chromosome 21q consisting of 591amino acids. It is mainly expressed in eukaryotes like humans and rat tissues. Being an organic secondary active anionic antiporter, it functions in uphill folate transport from blood into peripheral tissues by utilizing the high phosphate gradient. It is proved to be major facilitator system of folate in mammalian cell and tissue where it performs absorption process across intestinal and colonic epithelia. There are two more members of SLC19A1 i.e., SLC19A2 and A3 which functions in transportation in thiamine with 40% homology with A1. Folates have more affinity to transport through this transporter than folic acid and the activity of transportation is higher at neutral pH 7 but the activity constantly decreased below pH7. Human RFCs is N-glycosylated at Asn58 in loop domain connecting TMD1 and TMD2 Inactivation of RFCs in embryo is characterised by the folate deficiency and disorder. Structurally, three-dimensional structure of countable RFCs is available including bacterial (lacY) lactose/proton symporter. (Zhao, R., Matherly, L.H. and Goldman, I.D, Membrane transporters and folate homeostasis Expert reviews in molecular medicine, 11, 2009.)

## Material and method

### Identification of drug target

They are collectively an antiporter, symporter or uniporter system. They are found in different forms such as parasite Leishmania it is in form of FT1 and FT5 proteins, Slr0642 in Synechocystis and At2g32040 in Arabidopsis for transportation of folates in cells. The pteridine salvage by the help of biopterin transporters and folate transporters for the uptake of unconjugated biopterin and conjugated folate by the help of pteridine reductase enzyme. Which reduces oxidized biopterin to dehydration & tetrahydrate. Data provided the strong evidence for an unexpected complexity in mechanism employed to regulate pteridine at both RNA and protein levels (Mark et al 2000). Efforts to improve cancer and malaria treatment by modifying the genes playing role in folate pathway in mammalian cell which flashes a new role in drug discovery in mammalian cell (Nzila et al 2005). Inhibition of DHFR and DHPR was strategic for development of drug against malaria but reports suggested the findings of other folate pathway genes as a potential target. IC1D1964 was a potent inhibitor of mammalian thymidylate synthase inhibiting malaria weekly. The computational approach in comparing available genomic sequence of bacteria and Arabidopsis thaliana using SEED Database and its tools. The approach signifies the presence of novel GTP cyclopyrrolone and folyl polyglutamate synthase and p-amino benzoate gene whereas FolQ gene was reported to be missing gene from bacteria and plants. This led to the opening of many approaches for the genomic analysis of bacteria and plants (Laggard et al (2007). The prevalence global data for determination folate deficiency differing in different geographical distribution (Erin et al (2008). Data was collected from PubMed and vitamin and mineral nutrition information system at WHO. Out of 34 countries studied South-East Asia and Europe was found to be the most deficient countries in relation to folate. Folate assessment survey plasma serum concentration (55%), erythrocyte folate concentration (21%). No relation between vitamin concentration and geographical distribution was found.

The 3D structure of carboxyatractyloside inhibited ATP/ADP carrier. Bioinformatics studies suggest the study of 11 mitochondrial carriers, one of nuclear coded membrane proteins transporting variety of solutes, across mitochondrial membrane. Structure of carriers led to identification of different carrier in mitochondrial membrane based on knowledge of substrate specificity using homology modelling and multiple alignment analysis (Palmeiri et al (2009). The biology of different receptors of major facilitator family including (RFC), (PCFT), (FR) (Larry et al (2014). Reports suggested ubiquitous expression of RCFT in mammalian cell and tissue whereas PCFTs induce its expression in intestinal absorption and transportation in central nervous system. The major role of RFCs was reported to occur in transportation of antifolates like methotrexate and paratrexate, loss of which result into the resistance of methotrexate. Mechanistic aspect of folate regulation under folate deficient conditions. Results showed the intestinal uptake of folate by proton coupled folate transporter under acidic pH which showed the maximum transport by PCFT under acidic conditions (Wani et al (2012). mRNA expression profile analysis revealed the elevated increase in proteins level of both PCFT and RFC transporters suggesting the involvement of transcriptional and translational mechanism in regulating the intestinal folate uptake during folate deficiency.

Role of PCFT in folate deficient rats where upregulation folate under acidic pH increases by PCFT across intestinal brush border vesicle. The increment in folate uptake was associated with increase in V max with no change in Km of folate uptake process (Wani et al (2012). This suggested the upregulation of folate increases with folate deficiency which increases the activity of folate transporters but decreases the substrate binding like antifolates. RFC is reported to express in mammalian cell where it mediates the transport of one of the antifolate, methotrexate on the other hand PCFT is reported to transport the other antifolate, pemetrexed as well as intestinal transportation of folate. Both are excellent transporters but work in different ph. reviewed on the main methods used for the investigation of ion channel proteins. Molecular Dynamics is regarded as one of the best approaches to investigate the structure and dynamics in biological system. It also helps to understand the biophysical alterations induced by mutants (Boukhaba et al (2017). Molecular approach aims to characterize the ligand-protein and protein-protein interaction which allow to predict the optimal orientation between target proteins and molecules where each pose receive a score depending on the ligand and target fitting. Some of the descriptors like geometric, hydrophobic, lipophilicity and solubility are tabulated by QSAR modelling.

### Find Ligand for drug repurposing

**Table: - 1.** Drug Table.

| S. No | Candidate Drug molecules & Proton Pump Inhibitors | Compound ID (CID)<br>PubChem |
| --- | --- | --- |
| Substrate | Folic acid | Conformer3D_CID_135398658 |
| 1 | Myricetin | Conformer3D _ CID_ 5281672 |
| 2 | 2-[(4-Aminobenzoyl) amino]<br>pentanedioate-pABA-Glu | Conformer3D_CID_5103842 |
| 3 | 5-Methyl-THF (tetrahydrofolate) | Conformer3D_CID_135483998 |
| 4 | Methotrexate | Conformer3D_CID_126941 |
| 5 | Pemetrexed | Conformer3D_CID_135410875 |
| 6 | Raltitrexed | Conformer3D_CID_135400182 |
| 7 | 5H-Pyrrolo[3,2-d] pyrimidine | Conformer3D_CID_577022 |
| 8 | Aminopterin | Conformer3D_CID_169371 |
| 9 | Colchicine | Conformer3D_CID_6167 |
| 10 | Edatrexate | Conformer3D_CID_6917908 |
| 11 | Leucovorin | Conformer3D_CID_135403648 |

### Molecular Docking analysis

In this we used the MGL docking tool for the docking analysis. In this the grid box had formed that helped in the cover the target site. Retrieval of proteins sequence of hPCFT Uniprot/PDB. Protein sequence information from database will be used for Clustal Omega EBI search for evolutionary Phylogeny analysis. Secondary structure will be determined using I-TASSER and structure inspection by using Pymol. Preparation of library of potential repurposing inhibitors for hPCFT. Preparation of 3D structure library of potential repurposing inhibitors for hPCFT from PubChem. Molecular interaction analysis of individual drug inhibitor candidate against drug by Auto dock Vina, Auto Dock Tool (ADT) and PyMol.

### Screening of potential drug molecules to inhibit hPCFT

In recently hPCFT has been found to be inhibited by “myricetin” at a sustained mechanism (Yamashiro, T., et. Al, 2019). The Study of hPCFT myricetin sensitivity determine hPCFT segment involved in hPCFT – myricetin interaction revealed the segment from 83rd to 186th amino acid residue is directly involved in this interaction (Yamashiro, T., et. Al, 2019; Eudes, A., et al., 2010). Furthermore, the mutant G158 N-substituted hPCFT was found to be insensitive transformed to myricetin according to oppositely N158 G-substituted rPCFT was transformed to be sensitive to myricetin. Hence G158 of hPCFT acts as key residue for the These results indicates critical role of Gly158 in the myricetin sensitivity of hPCFT. Hence, we intend to screened potential drugs to inhibit the hPCFT on the basis of their stable molecular interaction with G158 residue of hPCFT.

### Other potential drugs to inhibit hPCFT

Pemetrexed is another excellent hPCFT substrate explain its demonstrated clinical efficacy for mesothelioma and non-small cell lung cancer, and prompted development of more PCFT selective tumor-targeted 6-substituted pyrrole pyrimidine antifolates that derive cytotoxic effects by targeting de novo purine nucleotide biosynthesis (Desmoulin, S.K., 2012). The Pemetrexed an antifolate that has been recently approved for treatment of mesothelioma and non-small cell lung cancer. DFD reduces MTD for methotrexate & raltitrexed ((located 50-fold), edatrexate (sevenfold), and pemetrexed (located 150-fold). Based upon lifespan extension, antitumor effect on methotrexate & edatrexate were greater in mice (L1210RFC) fed a folate-deficient diet (e.g., ILS: 455 and 544%, respectively) than in mice fed a standard diet (ILS: 213 and 263%, respectively). It was remarkably excellent. Regardless of dietary folate status raltitrexed and edatrexate were shown inactive in both mice containing L1210RFC and mice containing L1210MFR. This could be but due to high circulating plasma thymidine levels. Collectively, this study underscores that modulation of dietary folate status can provide a basis within which the therapeutic effect of antifolates may be further improved (Urquhart, B.L., et al., 2010). (hPCFT/SLC46A1) has been found to be recently potentially inhibit by myricetin. This raises a concern for malabsorption of folates in intestine. The colchicine, which is regarded as an inhibitor of vesicular-mediated endocytosis is observed to inhibit AP binding and AP-directed transport without affecting the basolateral transport.

### Potential drug molecules to inhibit hPCFT

Proton pump inhibitors are essential consideration group of drug candidates to screen antagonist for human PCFT. Hence the potential proton pump inhibitors are shortlisted and given here in table 2. These proton pump inhibitors are well known to inhibit proton pumps and are potential drug candidates for Folate pumps in different species. Hence, we utilized these shortlisted drug candidates to test for their potential to inhibit the active site of human PCFT.

**Table: - 2.** Drug Table Blocking amino acids of active site.

| S.No. | Drug Molecules | Binding Energy(kcal/mol) | Blocking amino acids of active site |
| --- | --- | --- | --- |
| **Substrate** | Folic acid | -6.8 | GLY-305, GLN-45, SER-83 |
| **1** | Myricetin | -7.6 | GLY-155, GLY-39, GLY156, PHE-157 |
| **2** | pentanedioate-pABA-Glu | -6.1 | GLY-158, LEU-36 |
| **3** | 5-Methyl-THF (tetrahydrofolate) | -5.9 | LYS-235, GLU-385, SER-172, LYS-381, SER-263 |
| **4** | Methotrexate | -6.7 | LYS-238, GLN-388, ASP-170, ARG-384, SER-172, ARG-177, HIS-499 |
| **5** | Pemetrexed | -7.8 | GLY-155, ALA-152 |
| **6** | Raltitrexed | -8.2 | THR-372, PHE-157, GLY157 |
| **7** | pyrimidine | -4.4 | GLY-155 |
| **8** | Aminopterin | -7.4 | GLY-155, PHE-157, SER-308, HIS-312 |
| **9** | Colchicine | -6.7 | ARG-22, ARG-14 |
| **10** | Edatrexate | -7.9 | GLY-155, PHE-157 |
| **11** | Leucovorin | -7.5 | GLY-97, GLY-93, THR-56, SER-83 |

### Raltitrexed and Edatrexate are the most potential drug against Cancer

The Binding energy of the Raltitrexed and Edatrexate was also comparatively lowest −8.2 and −7.9 kcal/mol respectively. These drugs cover the almost amino acids that is present on the protein (hPCFT) loop and block them active sites loop (THR-372, PHE-157,GLY157) and (GLY-155, PHE-157) of the molecules

## CONCLUSION

Folic acid transport is important for proper cell proliferation. In humans, folic acid is synthesized in body, so it is obtained externally. which are densely crowded in intestinal region. For the malignant cancerous cell these folates are crucially required on very frequent and enormous rate of fulfil the rapid proliferation of the cancerous cell. In the present study we have comparatively different PPI antagonist drug molecule which could be a potential candidate to inhibit the folate uptake by the (hPCFT) responsible to procure folate in abnormally enormous and frequent rate. These drug antagonist candidates finally include 27 as shortlisted based on previous studies. Out of these antagonist drugs, Raltitrexed and Edatrexate molecules were observed to specifically bind to the key active site loop (e.g., G155XXG158) of the hPCFT transporter. The Binding energy of the Raltitrexed and Edatrexate was also comparatively lowest −8.2 and −7.9 kcal/mol respectively. Hence from the present study we conclude that the Raltitrexed and Edatrexate antagonist drugs are the most potential inhibitory candidates for folate transport. We further tested the Raltitrexed drug candidate for its stable binding within the cavity of the hPCFT, and the results show potential inhibitory activity of the Raltitrexed drug candidate with stable binding to the hPCFT core binding site for a duration as long as 124 nanoseconds. Hence, overall, in the present study we may wind up that the Raltitrexed drug candidate would be a highly potential drug to inhibit folate up take through proton-pump hPCFT and hence the drug could be a potential anticancer drug to limit the proliferation of cancerous cells. Although in vivo and human trials would be an essential obligation to the said conclusions, which may be addressed in future prospects of the present study.

---

**Fig: 2.**
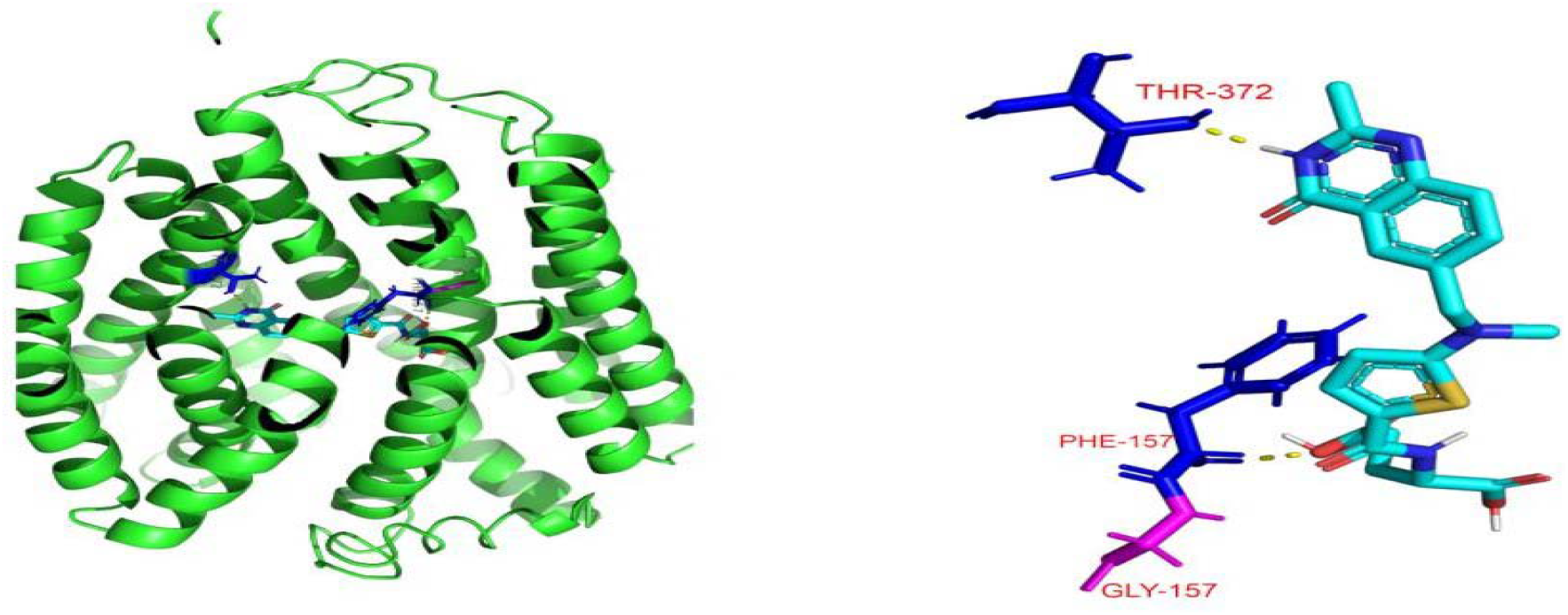
Our target site is GLY-156 in PCFT. Raltitrexed is binding with site GLN-157, PHE-157, THR-372 which is not far away from our target site and maybe can blocked the target site.

**Fig: 3.**
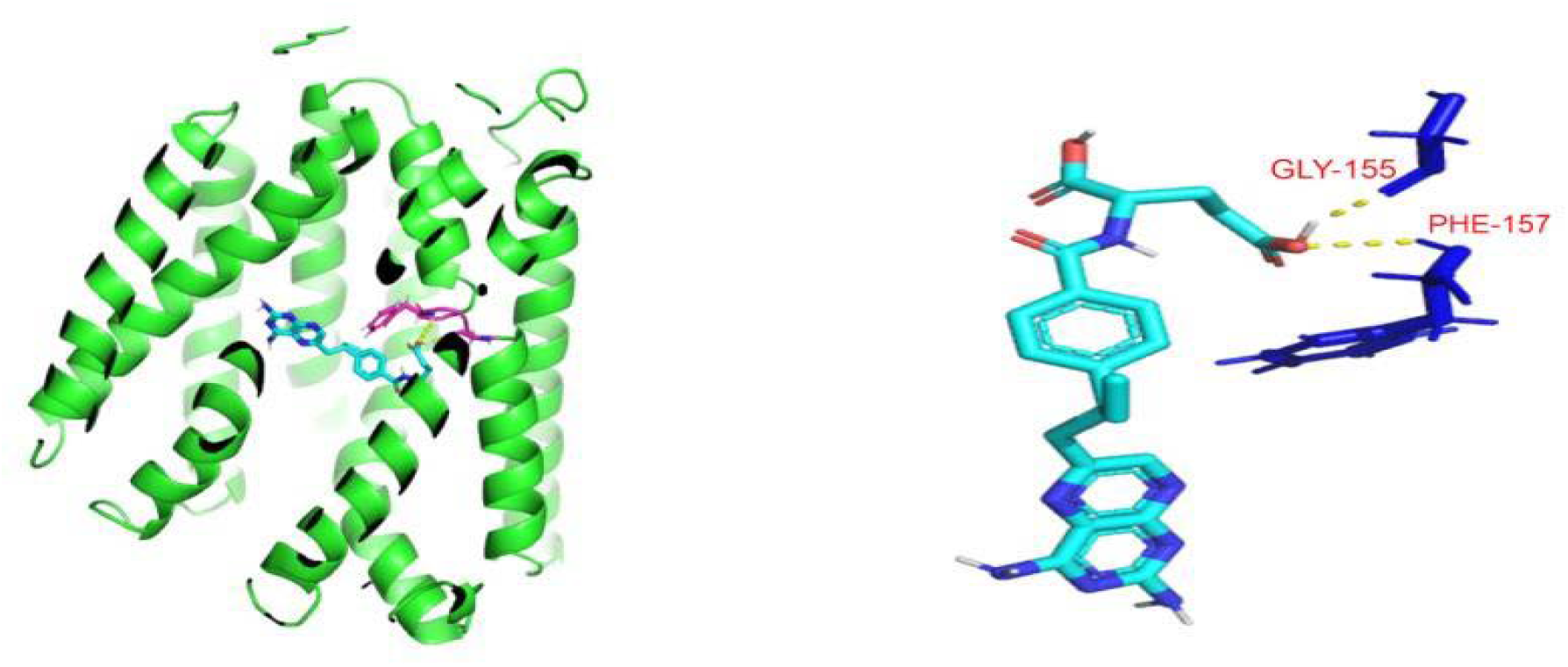
Our target site is GLY-156 in PCFT. Edatrexate is binding with site GLN-155, PHE-157 which is not far away from our target site and maybe can blocked the target site.

**Fig: 4.**
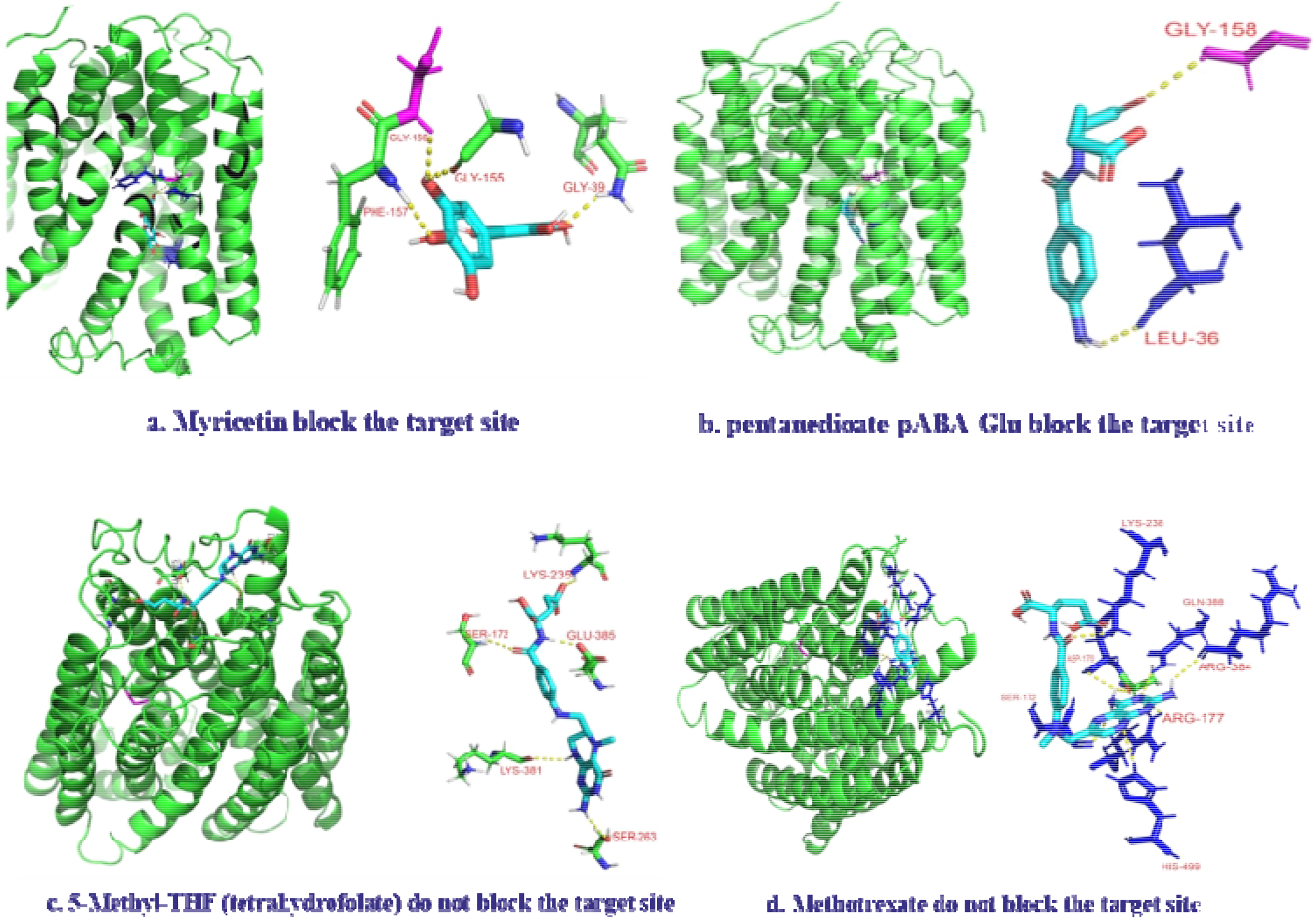
a. Myricetin is binding with site GLY-156, GLY-155, GLY-39, PHE-157. Myricetin is binding with target site and blocking the transportation of folate in human body. It has perfect credibility to repress the cancerous cell in human body but it has also negative effect on normal human health because it’s blocking the folate which is also required for normal day life. b. 2-[(4-Aminobenzoyl) amino] pentanedioate-pABA-Glu is binding with site GLY-158 and LEU-36 which is not our target site and not blocking the target site. c. 5-Methyl-THF (tetrahydrofolate) is binding withsite GLU-385, SER-172, LYS-235, LYS-381, SER-263 which is not our target site and not blocking the target site but covering whole protein. d. Methotrexate is binding with site GLN-388, SER-172, ASP-170, ARG-384, ARG-177, HIS-499, which is not our target site andnot blocking the target site but covering whole protein.

**Fig: 5.**
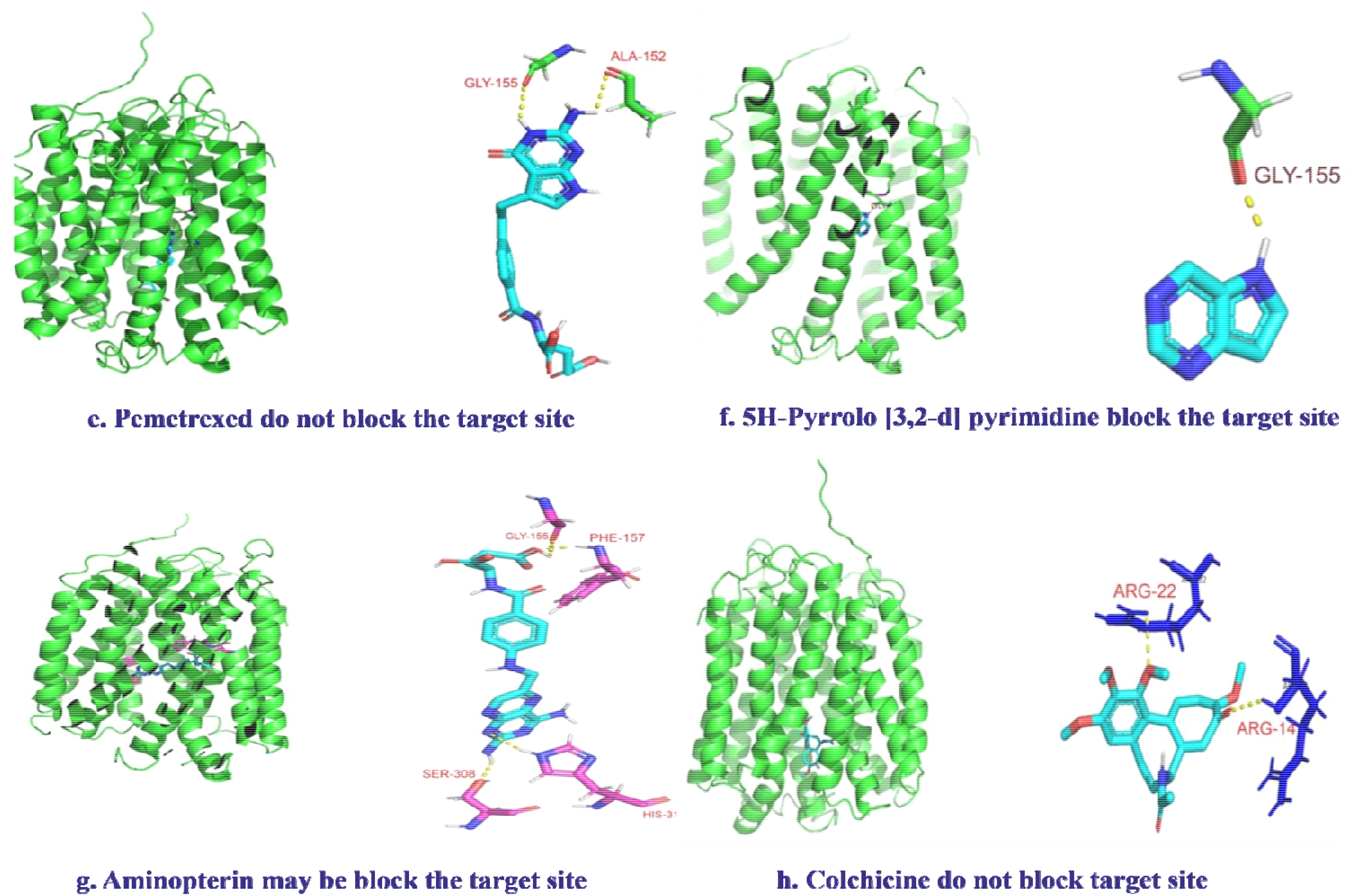
e. Pemetrexed is binding with site GLY-155, ALA-152 which is not far away from our target site maybe blocked the site. f. 5H-Pyrrolo[3,2-d] pyrimidine is binding with site GLY-155 which is not far away from our target site and maybe can blocked the target site. g. Aminopterin is binding with site GLY-155, PHE-157, SER-308, HIS-312 which is not far away from our target site and maybe can blocked the target site. h. COLCHICINE is binding with site ARG-22 and ARG-14 which Is not binding on our target site.

**Fig: 6.**
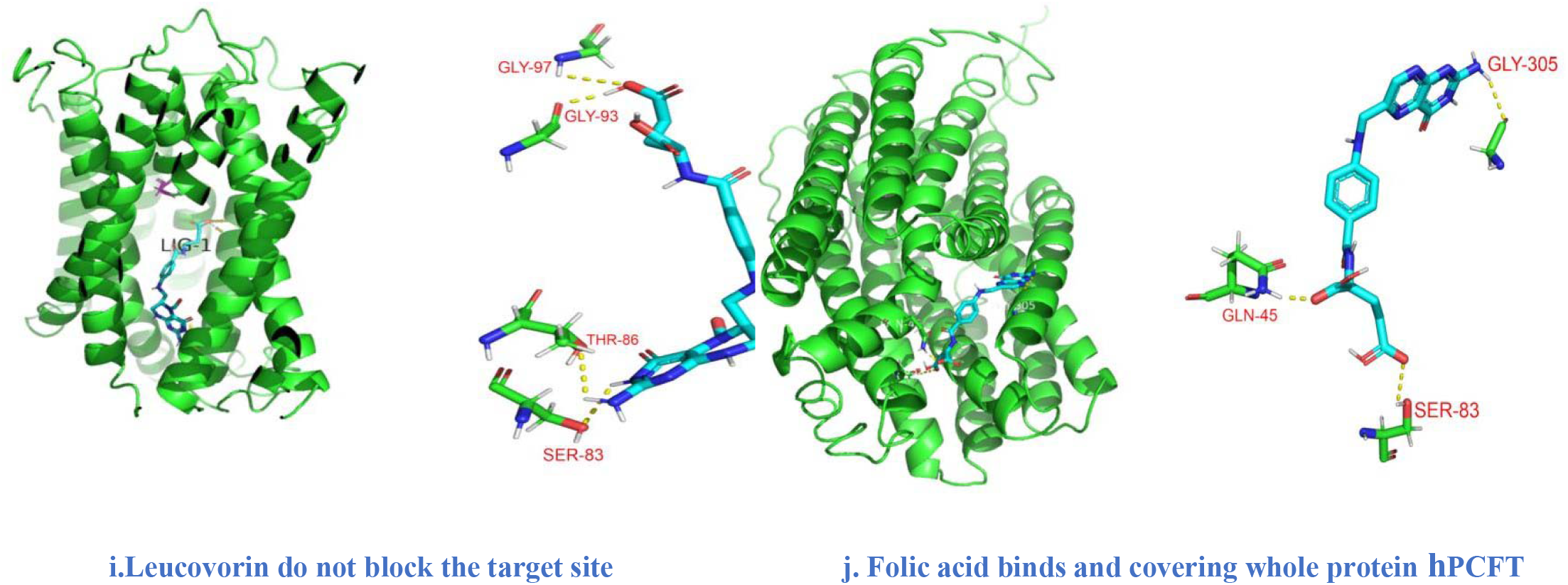
i. Our target site is GLY-156 in PCFT. Leucovorin is binding with site GLY-97, GLY-93, THR-86, SER-83 which is not far away from our target site and maybe can blocked the target site. j. Folic acid binds with sites GLY-305, GLN-45 and SER-83 and covering whole protein PCFT means folate transportation is blocked by folic acid by this cancerous cell in human body cannot absorb folate in rapid manner so it can help in reducing the generation of cancerous cell.

